# Local propagation dynamics of MEG interictal spikes: source reconstruction with traveling wave priors

**DOI:** 10.1101/2020.05.17.101121

**Authors:** Aleksandra Kuznetsova, Mikhail Lebedev, Alexei Ossadtchi

## Abstract

Epilepsy is one of the most common neurological disorders, with about 30% of cases being drug-resistant and requiring surgical intervention. To localize the epileptogenic zone (EZ), the pathological area that has to be surgically removed, brain regions are inspected for the presence of spikes during the interictal periods. This procedure maps irritative zones where spikes are present, but it is still challenging to determine which of the irritative zones generate seizures. To localize the source of seizures more precisely, a large-scale approach could be applied where the causal relationship is assessed between the signals recorded in a finite number of irritative zones [27]. This method however, does not reveal the fine-grained spatiotemporal patterns of spikes, which could provide valuable information regarding EZ location and increase the likelihood of surgery success [33].

Here we present a framework to noninvasively investigate the fine patterns of interictal spikes present in magnetoencephalographic (MEG) data. We use a traveling wave model, previously employed in the analysis of cortical alpha oscillations [16], to regularize the MEG inverse problem and to determine the cortical paths of spike traveling waves. Our algorithm represents spike propagation patterns as a superposition of local waves traveling along radial paths stemming from a single origin. With the help of the positively constrained LASSO technique we scan over wave onset moment and propagation velocity parameters to determine their combination that yields the best fit to the MEG sensor data of each spike.

We first used realistically simulated MEG data to validate the algorithm ability to successfully track interictal activity on a millimeter-millisecond scale. Next, we examined MEG data from three patients with drug-resistant epilepsy. Wave-like spike patterns with clear propagation dynamics were found in a fraction of spikes, whereas the other fraction could not be explained by the wave propagation model with a small number of propagation directions. Moreover, in agreement with the previous work [33], the spike waves with clear propagation dynamics exhibited spatial segregation and matched the clinical records on seizure onset zones (SOZs) available for two patients out of three.

## 1 Introduction

This study contributes to both the field of epilepsy research and the research on cortical traveling waves. Although cortical traveling waves have been known since the 1930s [1], [2], most studies of cortical activity have adhered to the Donders-paradigm assumption of space-time separability of brain activity [8] and relied on such neural data representation as across-trial averages – an approach that is unsuitable for propagating events with a high trial-to-trial variability [3], [4]. Yet, evidence of propagating patterns have been growing; they have been observed in many species and brain areas, including turtle visual cortex [29], visual, auditory and somatosensory cortices of the rabbit [12], sensorimotor cortex in awake mice [10], and primary and secondary visual cortices in awake monkeys [24]. Traveling waves have been found in human neocortex, as well (for a review see [23]), where, for example, alpha activity propagates [5], [16], [35] and exerts effects on gamma activity [5]). Additionally, theta traveling waves occur in both human neocortex [35] and hippocampus [18]. A traveling wave component is also present in the fixation-related lambda activity during free-viewing [13]. During sleep, large-scale [21] and local [15] propagation of slow waves takes place, and sleep-related K–complexes comprise intricate propagating patterns [19].

Although traveling waves appear to be ubiquitous in brain activity, their mechanisms and functions are far from being completely understood [9], [34]. Evidence, however, is mounting supporting two roles of traveling waves: (1) their functional role in normal brain processing, and (2) the contribution of propagating activity to brain pathological states. The normal functions of travelling waves are diverse. Thus, they contribute to working memory as suggested by the study where human subjects performed better on a memory task in the case their cortical propagating patterns were consistent [35]. Additionally, premovement and perimovement characteristics of cortical travelling waves correlate with the reaction time in a motor task, which indicates that traveling waves mediate motor control [28]. In the pathological domain, [17] described traveling waves in a Bicuculline model of epilepsy.

Here we examined traveling waves in the epileptic brain. Epilepsy is globally one of the most common neurological diseases, with 30% of cases being drug-resistant and needing surgical intervention. In a typical case of multifocal epilepsy, epileptic activity originates in the region called epileptogenic zone (EZ), or “seizure onset zone”, from where it propagates to the other parts of the brain, often covering the entire cortex and invading deep brain structures.

Identification of EZ is the main goal of presurgical diagnostics, where recordings of brain activity are examined during interictal and ictal periods to localize the site where epileptic activity starts. A large-scale approach is commonly used to tackle this issue, where the location of EZ is derived from the patterns of epileptic activity over the entire brain, see [27] for an attempt to formalize this. While this large-scale analysis is useful, the propagation dynamics of both interictal and ictal activity are multi-scale [30], [6]. At a macro scale, Tomlinson and his colleagues investigated interictal spike propagation using intracranial recordings with subdural grids and strips [33]. They suggested that the consistency of propagation direction could be used as a bio-marker of the epileptogenic region. Several studies examined local propagation of epileptic activity. Martinet and his colleagues [20] studied seizure dynamics on a millimeter spatial scale. Using 4*mm*^2^ microelectrode arrays, they showed that small groups of neurons spanning cortical columns generate rapidly propagating waves that could contribute to macro-scale seizures. Toward a better understanding of the local propagation of epileptic discharges, [7] developed a traveling wave model of ictal and interictal activity.

In this paper, we propose a method to explicitly localize traveling waves and determine their parameters from non-invasive magnetoencephalographic (MEG) recordings. We apply the proposed approach to analyse local propagation dynamics of the interictal spikes in patients with pharmacologically intractable epilepsy. MEG [25] has temporal resolution of several milliseconds and spatial resolution of several millimeters, which is suitable for analyzing local dynamics of the propagating epileptic activity. Moreover, one can gain insights on the anatomical paths of MEG travelling waves if appropriate methods are applied for solving the inverse problem for source localization. The problem of finding a unique solution to the inverse problem has fundamental limitations, so regularization methods are needed to deal with this issue [14]. To regularize the inverse problem, we model interictal spikes as waves travelling in radial directions from the source. This model works reasonably well both on simulated MEG data and the data from epileptic patients, where travelling-wave patterns were evident for a considerable fraction of interictal spikes stemming from a specific cortical region.

## 2 Methods

### 2.1 Data model

In our model, we consider an interictal spike as a traveling wave event. We suggest that the wave originates from a generating source and propagates in 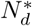 different directions along the cortical surface. While the distance covered by the wave depends on propagation speed, we set the length of all propagation paths equal in terms of the number of *N*_*s*_ cortical nodes visited. Thus, the *d*-th propagation direction path can be represented as a sequence of active cortical sources 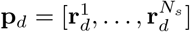, where **r**_*i*_ = [*x*_*i*_, *y*_*i*_, *z*_*i*_] contains source coordinates on a 3d-plane, 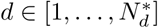, and the first source is the same for all directions (generator source).

The activation time series of the sources from the sequence **p**_*d*_ form the matrix **S**^*d*^ of size *N*_*s*_ × *T*_*s*_, where *T*_*s*_ = *T · fs* is the number of samples observed during the event, *T* is event duration in seconds, and *fs* is sampling frequency in Hz. In order to introduce the phenomenon of propagation in space and time, the source activation timeseries are shifted in time with respect to the activity of the preceding node.

Given the forward operator **G** with fixed source orientations and size *N*_*ch*_ × *N*_*src*_, where *N*_*ch*_ is the number of sensors and *N*_*src*_ is the total number of sources, multichannel MEG signals **X** can be represented as a linear combination of cortical traveling waves projected into the sensor space 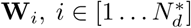:

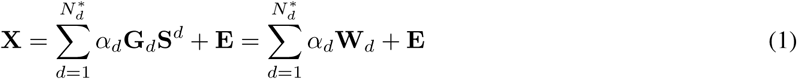

The matrix **G**_*d*_ of size *N*_*ch*_ × *N*_*s*_ is formed from the columns of the forward operator matrix **G** with topographies of the sources corresponding to the cortical mesh nodes along the path **p**_*d*_. The matrix **E** represents extraneous brain activity and additive sensor noise. The weighting coefficients 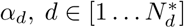 correspond to the contribution of each propagation direction to the observed MEG activity.

### 2.2 Basis waves

Given the data model introduced above, we suggest that propagating MEG activity can be represented as a linear combination of sensor-space traveling waves **W**_*i*_, *i* ∈ [1 *… N*_*d*_]. The main idea of the technique proposed in this paper is to generate templates of traveling waves, which we call *basis waves*, and then find their combination with the fewest number of terms that fits the MEG data. Below, we describe the algorithm for computing the basis waves.

As a simplification, we define the number of active cortical sources along each propagation path as equal to the number of observations made while the event lasts: *N*_*s*_ = *T · fs*. In our simulations we consider the case where the modeled activation time series for each of *N*_*s*_ sources have sinusoidal waveform and are shifted in time with respect to their sequence from the starting point.

For each propagation directions *d* ∈ [1, *…, N*_*d*_], source timeseries matrix **S**^*d*^ is formed of rows:

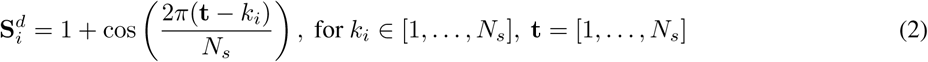

An example of a time series for *N*_*s*_ = 21 cortical sources located along a single propagation direction is shown in Figure 1. Each subplot corresponds to the activation profile of a single source *src*_*ind*_ ∈ [1, *…, N*_*s*_]. Maximal activation amplitude time points are marked with red dots for each source. The choice of this time series function allows us to model the propagation of activity in both space and time.

**Figure 1:**
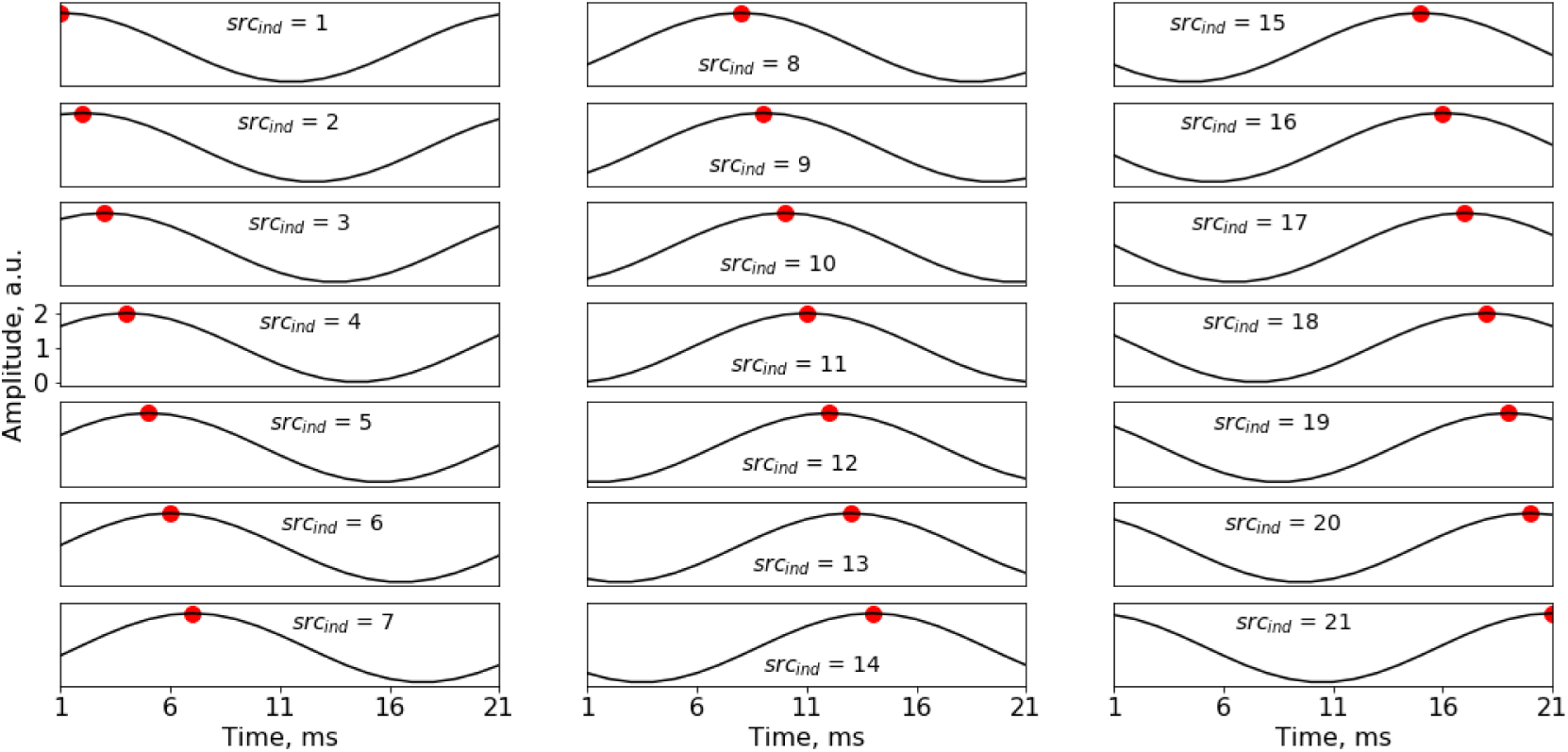
Activation timeseries for *N*_*s*_ = 21 sources, modeling the propagation of maximal activation (red dot) both in space and time.

Source locations 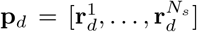 in each concrete case depend on the individual anatomy, the starting source **v**_*s*_ = [*x*_*s*_, *y*_*s*_, *z*_*s*_] and the propagation velocity. Cortical paths for the basis waves are generated using Freesurfer surfaces [11] computed in Brainstorm software [31]. Thus, for each basis wave, we need to define the path in the graph with *N*_*src*_ vertices connected according to the adjacency matrix **A** defined by the cortical model. For a given starting cortical location with *N*_*d*_ nearest neighbours, we define *N*_*d*_ basis waves propagating in the nearest neighbour directions. To make the analysis procedure convenient for practical applications, we do not add new vertices or edges to the cortical model graph. A limitation of this approach is that the number of propagation directions depends on the density of the vertices in the region of interest and in case of adaptive meshes, on its curvature. This makes sense since the spatial resolution of MEG is positively correlated with the local curvature.

Propagation paths for starting location **v**_*s*_ are generated according to the following algorithm:

1. The *N*_*d*_ nearest neighbour vertices 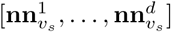 for a starting point are found in the corresponding row of adjacency matrix 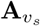.
2. To approximate the vector normal to a cortical patch formed with nearest neighbour vertices 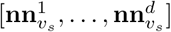, we average the normal vectors in each of *N*_*d*_ locations: 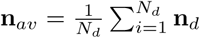. The unit-norm version of this vector is 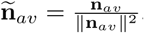.
3. Calculate the matrix projecting into this cortical patch 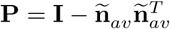, where **I** is the 3 × 3 identity matrix and (*·*)^*T*^ is the matrix transpose operator.
4. 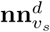 is added to the path **p**_*d*_
5. The propagation direction vector is computed: **h**_*sd*_ = (**v**_*s*_ − **v**_*d*_) *·* **P**. The normalized version of this vector is 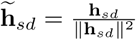.
6. For each nearest neighbour, while the path is shorter than *max*_*step*, the following operations are repeated:
  a. Assign the nearest neighbour vertex to auxiliary variable **v** and using the adjacency matrix find its nearest neighbours [**nn**_1_, *…*, **nn**_*m*_]
  b. Among all vertices found choose the one maximizing the criterion 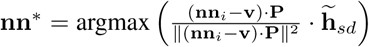
  c. Add the vertex **nn**^∗^ to the path **p**_*d*_ and repeat step 6.

The obtained cortical paths **t**_*d*_ are then used to define the particular source locations (nodes) **p**_*d*_ for different propagation velocities. Figure 2 shows an example of the generated sets of sources **p**_*d*_ for different propagation velocities, from 0.3 m/s to 1.5 m/s. Given the propagation paths, the forward operator matrix subsets **G**_*d*_, *d* = 1 *… N*_*d*_ are then used to calculate the basis waves following equation (1).

**Figure 2:**
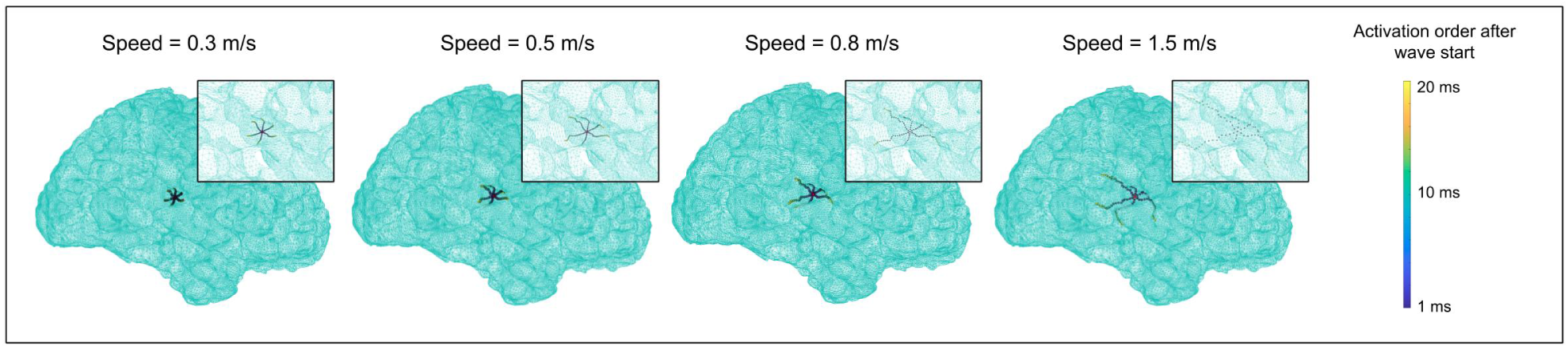
Example of generated basis waves source locations for *N* = 6 and different propagation velocities

In addition to the plane-directed waves, we also consider a spherical wave propagating in all directions simultaneously and composed of a sum of plane waves 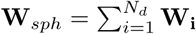. However, our tests on simulated and real data have shown that spherical waves are not selected by the algorithm as members of the optimal combination.

When varying the propagation velocity value, we also introduce a timestamp of wave initiation. The exact time of wave initiation is unknown, but an optimal value can be found using a sliding window. We automatically scan the time interval containing an interictal spike, fit the basis waves to this interval and then repeat the whole analysis for the time series delayed by one sample.

### 2.3 Best combination of traveling waves

After the basis waves have been generated, the next step of the analysis consists of searching for their combination that best describes the observed MEG data. Based on physiological assumptions, the desired combination should contain only several basis waves corresponding to several dominant propagation directions. Therefore, we are looking for the most sparse solution that describes the data which corresponds to a small count of well defined dominant propagation directions.

We used LASSO technique [32] to find the contribution of each precomputed basis wave to the MEG data, with an additional constraint that LASSO coefficients were positive. The optimization problem is formulated by equation (3). The main advantage of this technique is that, due to the non-smooth *L*_1_ regularization term, it allows to perform feature selection in such a way that the coefficients of non-informative propagation directions are equal to zero.

To address the multi-channel problem, we vectorize the data matrix **X** and basis waves on sensors 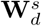.

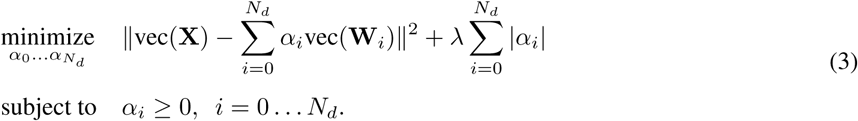

This procedure is then applied to all sets of generated basis waves with two parameters: the propagation velocity and wave onset time. The best solution is selected according to *R*-squared (i.e. ratio of variance explained).

The important issue in basis waves generation is the detection of the very first source that initiates the propagating pattern. We define the region of interest (ROI) in the first approximation with the RAP-MUSIC dipole fitting algorithm [22]. In order to improve the solution accuracy, we scan the ROI using the vertexes as starting points and compare the solutions using the *R*^2^ metrics.

### 2.4 Algorithm pipeline

The final pipeline for solving MEG inverse problem with traveling wave priors comprises the following steps:

1. Determine the cortical ROI by applying to the MEG data the RAP-MUSIC algorithm [22] and picking its vicinity (for example, 1 cm radius). Then for each vertex that belongs to this ROI, repeat:
  a. Use the picked cortical source as the starting point for wave generation **v**_*s*_ and calculate the basis waves for different propagation velocities.
  b. Use the LASSO technique [32] with positive coefficients to fit the basis waves to the very beginning of MEG recording of the considered event and determine the optimal *R*^2^ value and the number of nonzero coefficients in the solution.
  c. Delay the MEG signal by *p* samples and repeat the analysis; then find the optimal starting timestamp and the optimal propagation velocity according to the *R*^2^ metric.
2. Compare optimal *R*^2^ for all vertices in the ROI and pick the best generation point as the closest to the real source.

### 2.5 Monte Carlo simulations

In order to test the algorithm performance, we performed a set of Monte Carlo simulations. Synthetic MEG magnetometer data was generated with a high-resolution cortical surface model with 300,000 vertices reconstructed from anatomical MRI data using FreeSurfer software [11]. The forward-model matrix **G** for dipolar sources with fixed orientation was computed using Brainstorm software [31] with the overlapping spheres technique.

The propagation speed considered in the analyses of both simulated and real data was taken from the following set: **S** = [0.001, 0.005, 0.01, 0.05, 0.1, 0.2, 0.3, 0.4, 0.5, 0.6, 0.7, 0.8] m/s. We analyzed 300 trials with a uniformly distributed propagation speed (25 trials per each value). Wave duration was set to *T* = 20 ms, and the sampling frequency was 1,000 Hz.

In simulations we considered two types of trials. The first was a simulation of a traveling wave (300 trials), and the second was a simulation of non-propagating oscillatory activity (an oscillating blob; also 300 trials). Traveling waves were generated with the algorithm described above in section 2.2. The oscillating blob was computed as 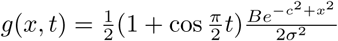.

We defined the SNR for the simulated data in the sensor space as the ratio of Frobenius norm for 5 channels with the highest power during the spike interval to the norm for the data from the same channels during the period of the same duration preceding or succeeding the interictal spike. We assured that the latter period did not contain spikes.

For each type of trial, a randomly picked vertex played the role of a generating source. The propagation direction and speed were randomly picked from the available set of parameters.

We modeled extraneous activity with *Q* = 1, 000 task-unrelated cerebral sources whose locations and time series varied across epochs. The timeseries of the task unrelated sources were narrow-band signals obtained by means of zero-phase filtering of the realizations of Gaussian pseudorandom process. The filtering was performed with the fifth-order band-pass IIR filters; the filter bands corresponded to theta (4-7 Hz), alpha (8-12 Hz), beta (15-30 Hz) and gamma (30-50 Hz, 50-70 Hz) activity. We adjusted the relative contributions of these rhythmic components according to the well-known 1*/f* characteristic of the MEG noise spectrum. The noise components corresponded to the typical signal-to-noise ratio of MEG recordings. These sources were projected into the sensor space using the corresponding columns of the forward matrix. We simulated 200 epochs of MEG data. For each epoch, a new randomly picked set of noisy sources was chosen, and new noisy time series were generated with a 1*/f* spectrum.

The high-resolution grid was used only for data simulation. For source reconstruction, we employed a sparser cortical grid with 200,000 vertices. Additionally, we artificially introduced an error for detecting the first source. To do this, we first randomly picked a cortical generating location from the dense model and then used this location as a starting point for the wave. Next, we randomly derived the estimated starting point using the sparse model for the 3-mm area around the exact generating location. The basis waves were initiated at this new location.

To compare the generated and estimated propagation direction, we calculated the first principal direction **p**^∗^ for the ground truth direction. We then determined all directions used in the estimated solution and assigned them the weights 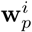 that were proportional to their contribution to the LASSO solution. We then calculated the weighted first principal propagation direction for the optimal solution 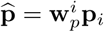. The direction estimation error was calculated as 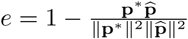.

To minimize the wave detection errors due to an inaccurate first point localization, we scanned the 5 mm area around those points and picked the best generating point as the one with the lowest *R*^2^.

### 2.6 MEG data acquisition

We applied the proposed algorithm to the MEG datasets from three patients with multifocal epilepsy. The data were collected at Moscow MEG facility with Elekta-Neuromag Vectorview 306 system (Elekta Oy, Finland) that had 204 planar gradiometers and 102 magnetometers. The data were collected during sleep at the sampling rate of 1000 Hz, and preprocessed using Elekta MaxFilter™ software.

## 3 Results

### 3.1 Monte Carlo simulation

Monte Carlo simulations were performed for three SNR levels: SNR equal to 1, 2, and 3. These results are shown in the top left panel of Figure 3 where the receiver operating characteristics (ROC) curves are depicted for wave detection. The detection was tested on 300 Monte Carlo trials with having an uniform distribution of propagation speed and 300 trials with non-propagating activity. The corresponding AUCs are 0.78, 0.95, and 0.97, indicating that the proposed technique successfully discriminated between the propagating and non-propagating activity when SNR was reasonably high.

**Figure 3:**
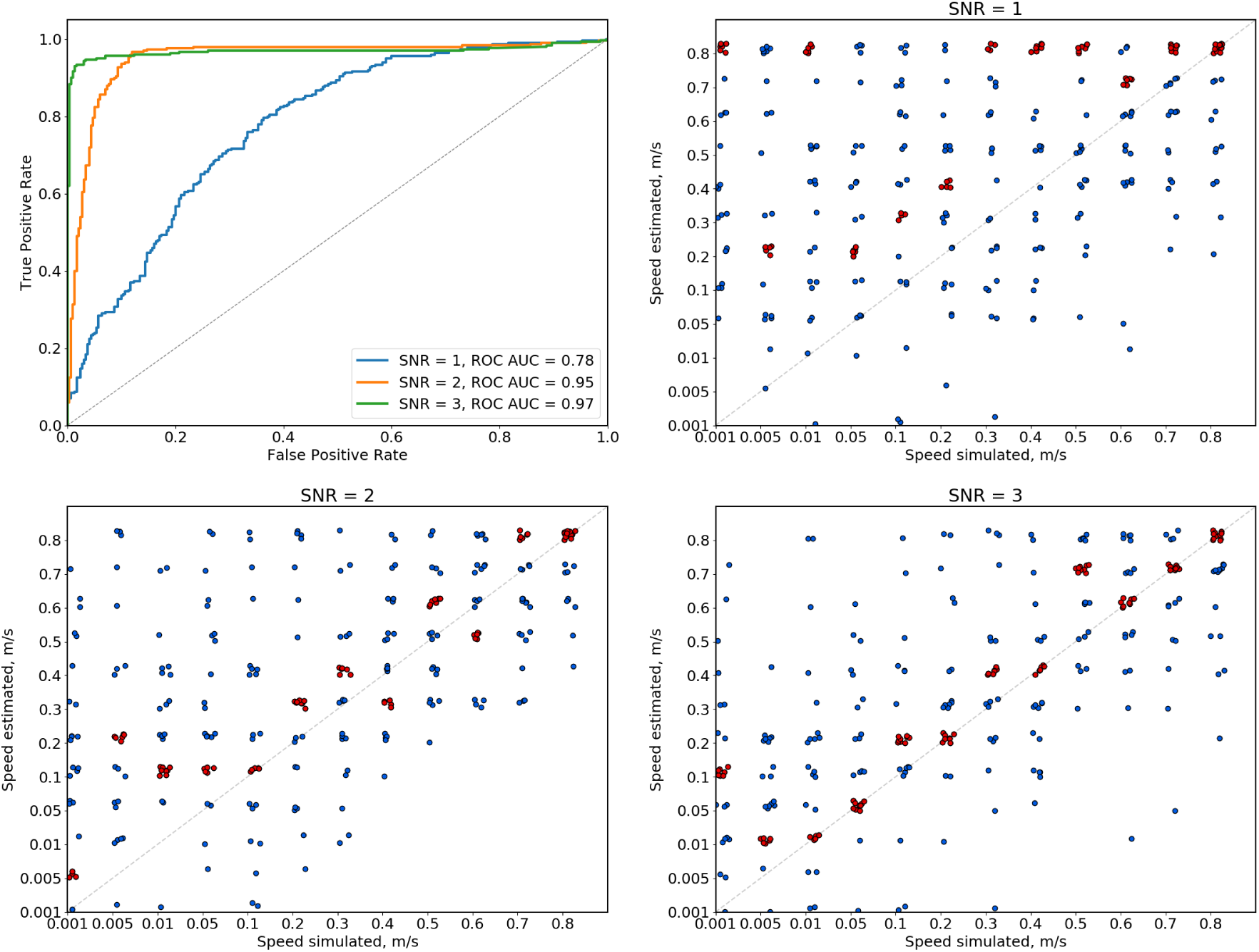
Comparison of simulated (x-axis) and estimated (y-axis) speeds for three SNR levels: SNR equal to 1, 2, or 3. Each dot corresponds to one Monte Carlo trial. A small random jitter was added to the values for visualization purposes. The most frequent estimated speed is determined for each ground truth speed value, and the corresponding dots are shown in red.

The other panels of Figure 3 compare the simulated (x-axis) and estimated (y-axis) speed values for different SNRs. Each dot corresponds to one Monte Carlo trial. A small random jitter was added to the values for visualization purpose. The most frequent estimated speed values were determined for each ground truth speed and plotted as red dots. For SNR equal to 1, the algorithm tended to significantly overestimate the propagation speed as compared to the ground truth value: the clusters of red dots do not coincide with the ground truth except for the highest propagating velocity values. For SNR equal to 2, erroneous detections are still plentiful, but the absolute difference between the estimated and the actual propagation speed is much lower than for SNR of 1. For SNR equal to 3, the most frequent value of the estimated speed coincides with the actual speed or with the closest value for all speeds except for two cases, where speed is overestimated.

It is important to note, that errors in estimating speed are inevitable even for SNR because of the error in the starting point localization and the usage of a sparse model. If the estimated starting point of a wave is shifted relative to the actual starting point toward the end of the propagation path, speed is underestimated and, conversely, if the starting point is shifted away from the path endpoint speed is overestimated. The higher is the SNR, the lower are these errors.

Next, we evaluated the errors in the estimated propagation direction. Figure 4 shows the distribution of these errors for the three SNR levels. The error was calculated as 1 − cos(*ϕ*), where *ϕ* is the angle between the actual and estimated principal direction. The error ranges between zero and one. For all SNR levels, the error is typically less than 0.1 and it decreases with the the increase in SNR.

**Figure 4:**
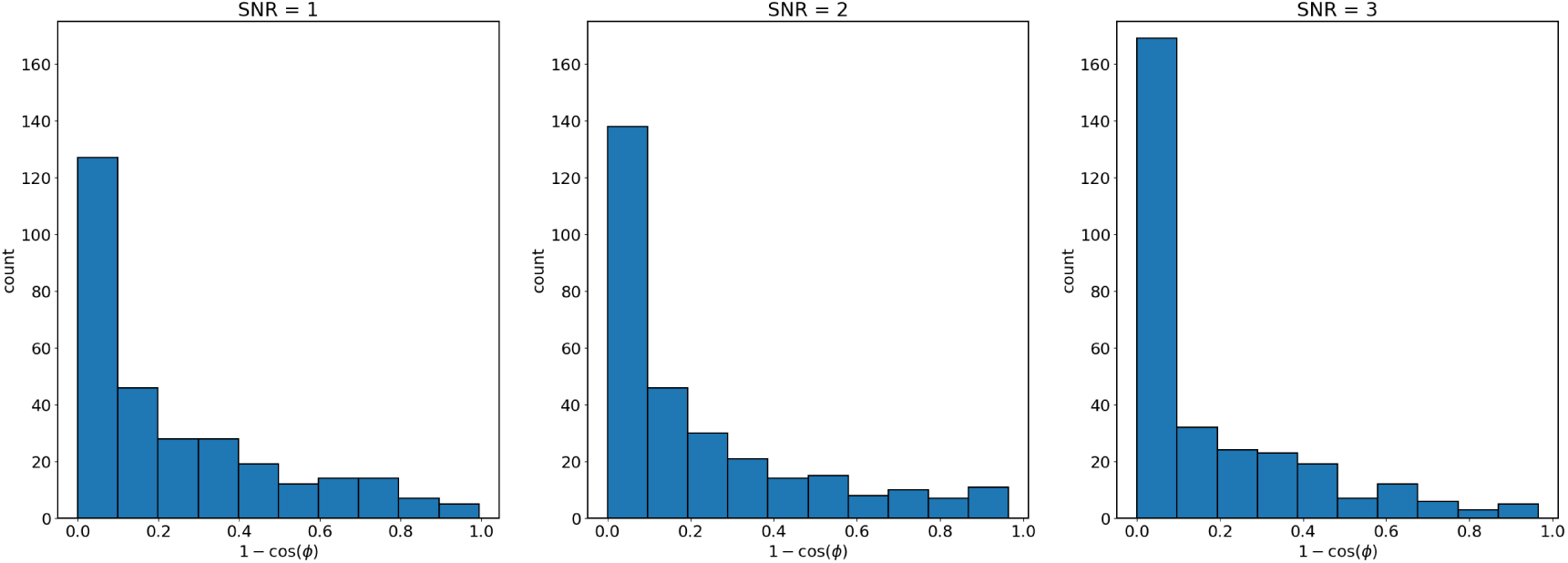
Statistical distributions of the error in the estimation of propagation direction, 1 − cos(*ϕ*), where *ϕ* is the angle between the principal directions of the actual and estimated propagation. The results are shown for 300 Monte Carlo trials and for three SNR levels: SNR equal to 1, 2, or 3.

### 3.2 Patient data

The data from three patients consisted of 10-min MEG recordings conducted during sleep. We detected interictal spikes in these data using the ASPIRE technique [26], which was based on the independent component analysis (ICA) decomposition. We started with selecting the independent components with the most clear spike patterns. This procedure was performed as ICA decomposition with the Infomax technique. A simple threshold was then applied to detect the occurrences of spikes. Next, we fitted dipoles with the RAP-MUSIC algorithm [22] to localize the spike-generating sources and identify the physiologically plausible detected events. We used 0.97 as the threshold for the subspace correlation metrics. Finally, we applied a simple deterministic distance-based clustering algorithm to combined the sources into tight clusters with a radius of 1 cm, each containing ten or more dipoles. The ASPIRE parameters were found empirically; they were set to be identical for all patients. Although the detection procedure was run separately for the gradiometers and magnetometers, roughly the same clusters were found. All the subsequent analyses utilized MaxFiltered magnetometer data.

#### 3.2.1 Analysis of individual spikes

The algorithm proposed here was applied to each interictal spike separately. Figure 5 illustrates a detailed analysis of a single spike. The starting point is determined using a sliding window method, where basis waves have a duration of 20 ms, and the duration of MEG signals is 40 ms. Panel A of the figure shows *R*^2^ values obtained for different propagation velocities (x-axis) and starting timepoints (y-axis). Panel B depicts the number of nonzero coefficients in the solution. The best solution is determined as the one with the highest *R*^2^. The *R*^2^ values color-plotted as a function of the starting point and propagation speed reveal a clear peak, marked with the red dot. For this particular spike, the highest *R*^2^ is 0.72, which corresponds to a physiologically plausible propagation speed of approximately 0.3 m/s. A single propagation direction is sufficient to characterize this spike. Panel C shows the spike time course, with the time interval best explained by the travelling-wave model highlighted. Note that the traveling wave model is in a particularly good correspondence with the upward-slope portion of the spike.

**Figure 5:**
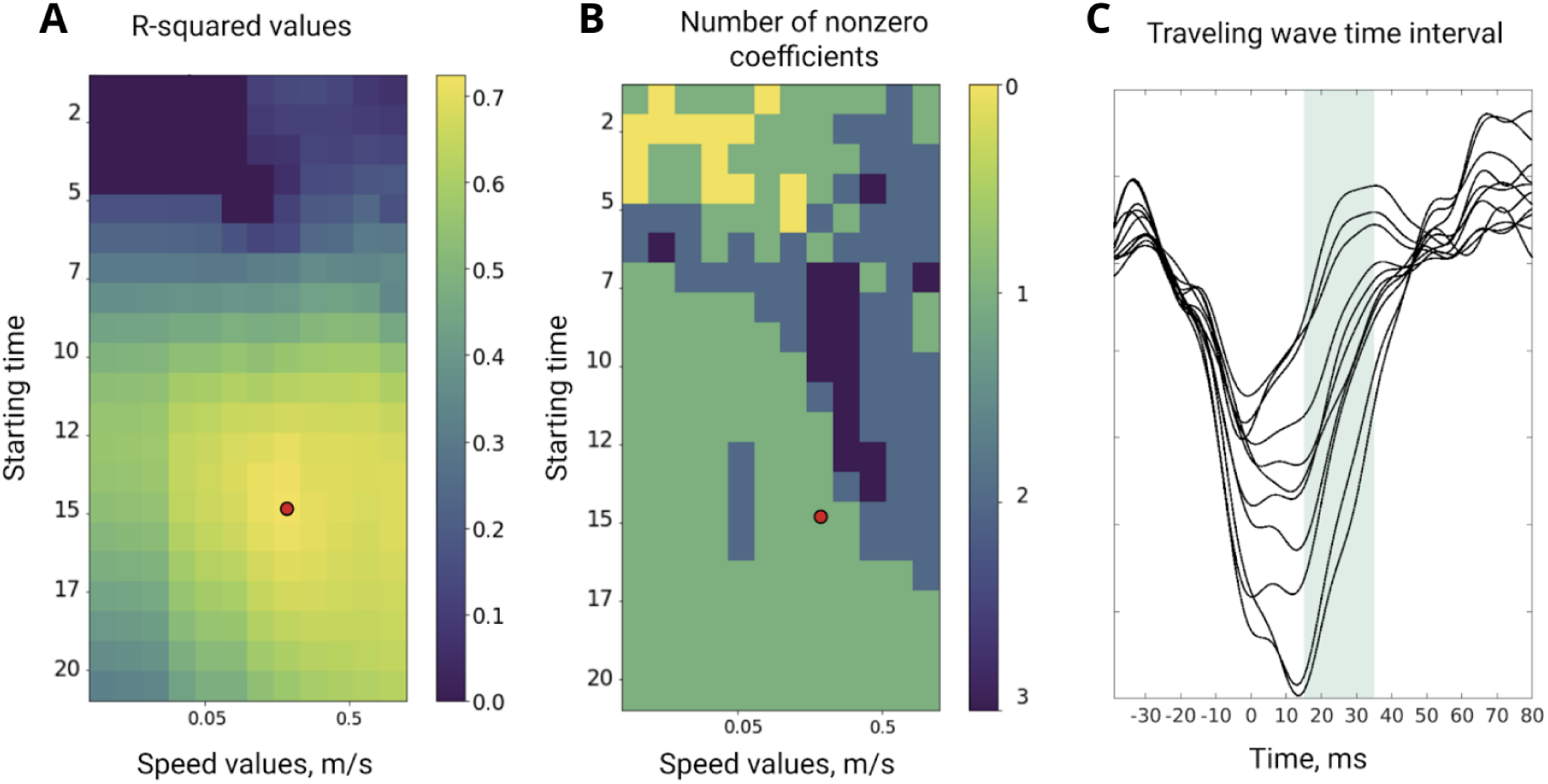
A representative example of a single interictal spike analysis. **A**. *R*^2^ values obtained for different starting time points and propagation speed values with the optimal solution marked in red. **B**. Corresponding numbers of nonzero coefficients. **C**. MEG timeseries of the considered spike, best fitted time interval highlighted.

#### 3.2.2 Aggregated results for three patients

The descriptive statistics for all patients are shown in Table 1. The variable *N*_*spikes*_ is the number of spike events in each particular cluster, *T*_*start*_ is the time of the first event from a specific group, measured in seconds from the beginning of the recording, and *T*_*var*_ is the standard deviation of event timepoints in a cluster, measured in seconds.

**Table 1:** Descriptive statistics for all clusters for three patients.

| Cl | Patient 1 |  |  | Patient 2 |  |  | Patient 3 |  |  |
| --- | --- | --- | --- | --- | --- | --- | --- | --- | --- |
| | $N_{spikes}$ | $T_{start}$ | $T_{var}$ | $N_{spikes}$ | $T_{start}$ | $T_{var}$ | $N_{spikes}$ | $T_{start}$ | $T_{var}$ |
| 1 | 64 | 10.570 | 99.005 | 233 | 44.728 | 140.120 | 114 | 2.348 | 99.461 |
| 2 | 18 | 9.349 | 122.090 | 10 | 56.844 | 163.160 | 34 | 77.175 | 136.020 |
| 3 | 27 | 16.372 | 85.610 | 52 | 120.883 | 144.150 | 14 | 141.136 | 134.140 |
| 4 | 12 | 10.387 | 128.120 | 21 | 56.789 | 146.560 |  |  |  |
| 5 | 38 | 12.556 | 99.757 | 19 | 176.413 | 134.220 |  |  |  |
| 6 | 13 | 27.265 | 85.078 | 12 | 151.483 | 151.270 |  |  |  |
| 7 | 29 | 2.442 | 110.200 | 13 | 59.603 | 95.913 |  |  |  |
| 8 |  |  |  | 9 | 63.843 | 174.020 |  |  |  |
| 9 |  |  |  | 9 | 1.001 | 123.360 |  |  |  |
| 10 |  |  |  | 10 | 82.998 | 118.340 |  |  |  |

We applied our technique to each detected interictal spike and aggregated the obtained *R*^2^ based on their assignment to the clusters. Since the goal of this analysis is to find the parsimonious description of an interictal spike, the other critical factor is the number of propagation directions for the optimal solution. Figure 6 demonstrates the location of seven epileptic foci (shown with different colors) and the distributions of the two chosen metrics for Patient 1. The left panel for each cluster shows the distribution of *R*^2^ values for the spikes forming this cluster; this distribution characterizes the traveling wave model goodness of fit. The right panel for each cluster shows the number of nonzero coefficients, which characterizes the model simplicity.

**Figure 6:**
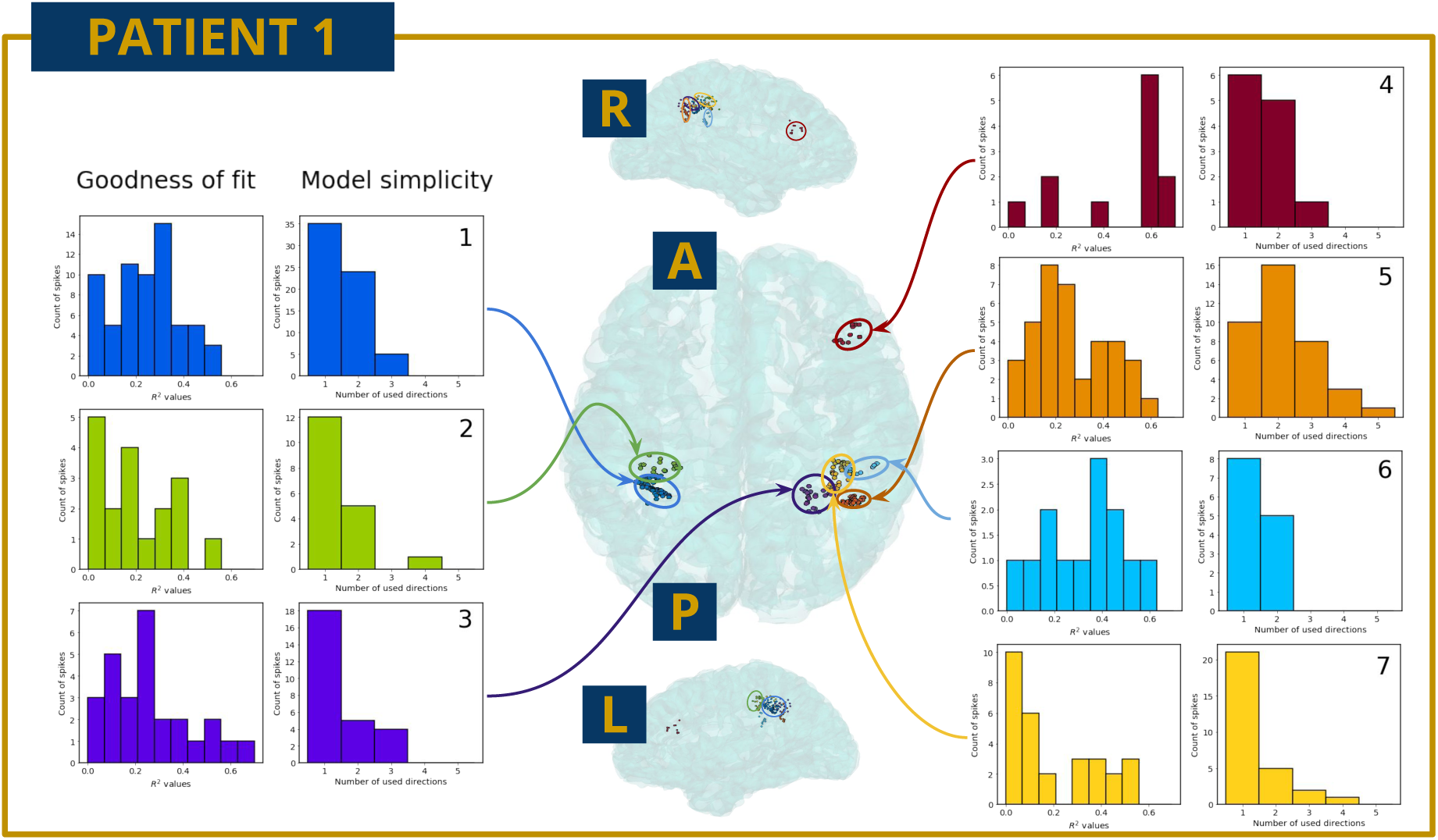
The localization of seven epileptic foci automatically detected with the ASPIRE technique for Patient 1: top axial view in the middle and right and left hemisphere sagittal view on the top and the bottom correspondingly. The distribution of *R*^2^ metrics (goodness of fit) and number of direction used in the optimal solution (model simplicity) for each cluster.

Figure 7 shows the analysis results for the Patient 2 and Figure 8 for the Patient 3.

**Figure 7:**
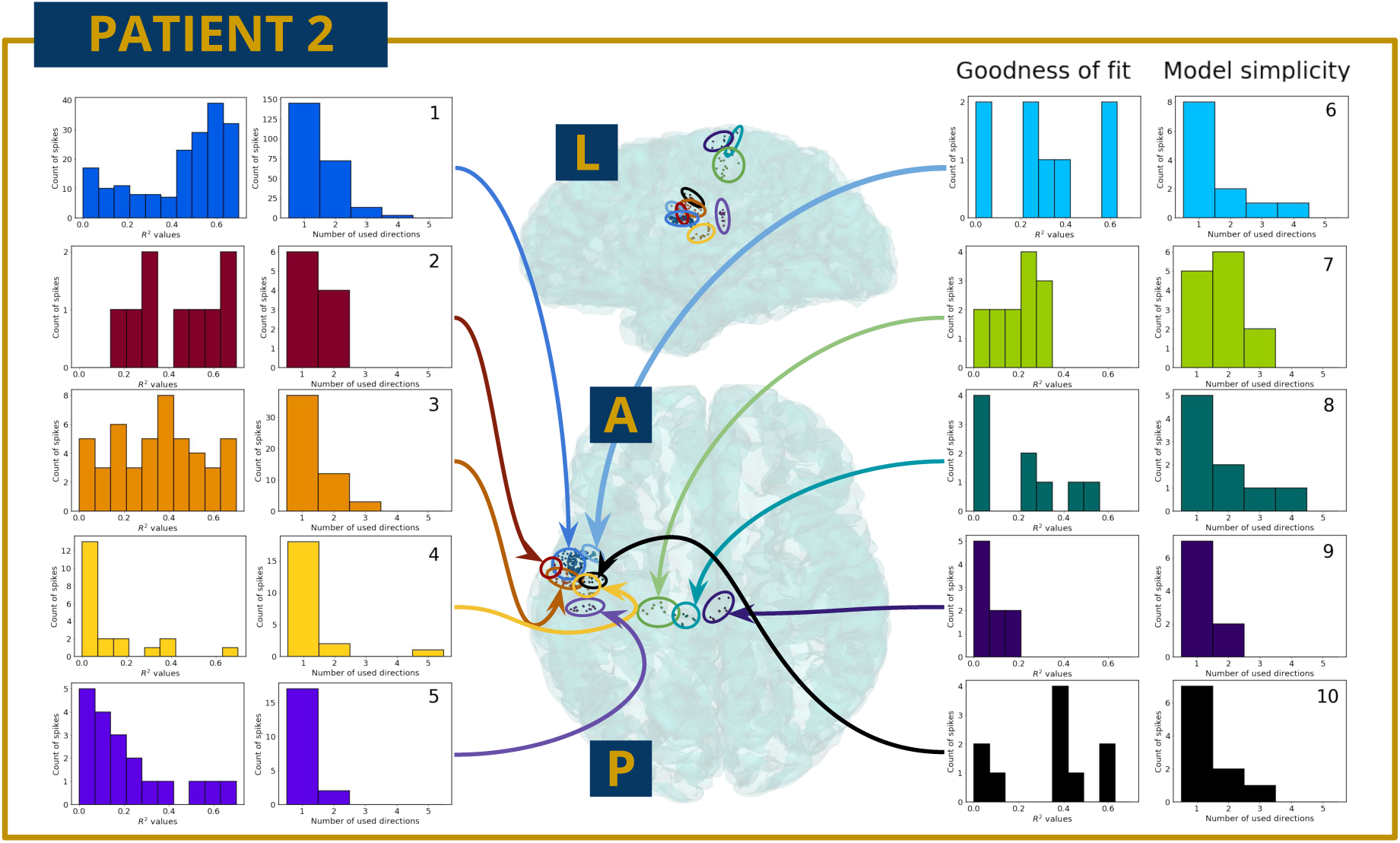
The localization of ten epileptic foci automatically detected with the ASPIRE technique for Patient 2: top axial view in the middle and left hemisphere sagittal view on the top. The distribution of *R*^2^ metrics (goodness of fit) and number of direction used in the optimal solution (model simplicity) for each cluster.

**Figure 8:**
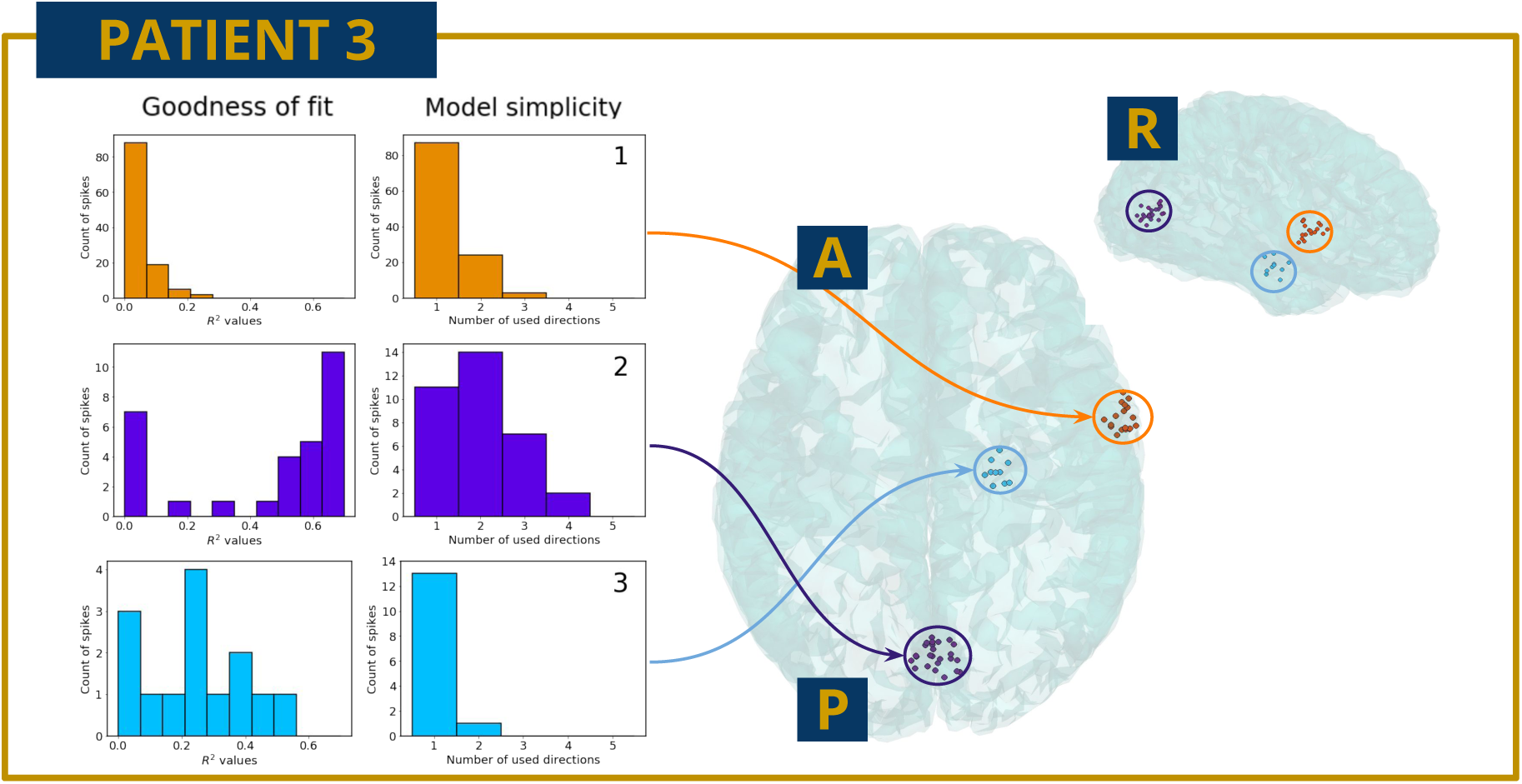
The localization of three epileptic foci automatically detected with the ASPIRE technique for Patient 3: top axial view and right hemisphere sagittal view. The distribution of *R*^2^ metrics (goodness of fit) and number of direction used in the optimal solution (model simplicity) for each cluster.

The analysis of patient data revealed a variability of the wave-model goodness of fit depending on the concrete spikes being examined. A parsimonious wave propagation model with a few dominant directions fit only a portion of the analyzed spikes. Table 2 shows the proportion of spikes with the goodness of fit higher or equal to 0.6 for each cluster and for all patients. For Patient 1, only Cluster 4 data is in good correspondence with the travelling-wave model as it contains 67% of well-fitted spikes. For the other two patients, the percentage of spikes explained by the wave propagation model varies significantly between clusters. Cluster 1 of Patient 2 contains the largest proportion of wave-like spikes, followed by Cluster 2 whose spatial characteristics are close to the ones of Cluster 1. The other clusters contain a few of wave-like events. For Patient 3, one occipital cluster shows a large portion of spikes with good correspondence to the model, whereas the others do not match the model well.

**Table 2:** The percentage of spikes well-explained with a traveling wave model in each cluster in three patients. We consider an interictal spike as well-explained, if *R*^2^ is greater or equal to 0.6.

|  | Patient 1 | Patient 2 | Patient 3 |
| --- | --- | --- | --- |
| 1 | 3 % | 70 % | 0 % |
| 2 | 5 % | 57 % | 69 % |
| 3 | 14 % | 34 % | 7 % |
| 4 | 67 % | 4 % |  |
| 5 | 10 % | 11 % |  |
| 6 | 15 % | 25 % |  |
| 7 | 10 % | 0 % |  |
| 8 |  | 11 % |  |
| 9 |  | 0 % |  |
| 10 |  | 30 % |  |

For the simulated data, the goodness of fit depends on spike SNR, as shown in Figure 3. Therefore, SNRs of individual spikes could explain why some of the spikes, recorded in patients, were well explained by the wave-propagation model and the others were not. To clarify this issue, we analyzed the distribution of *R*^2^ as a function of the SNR of individual spikes. The results of this analysis for all three patients are shown in Figure 9. According to this analysis, there is no significant dependence of the goodness of fit on the SNR in any of the three patients.

**Figure 9:**
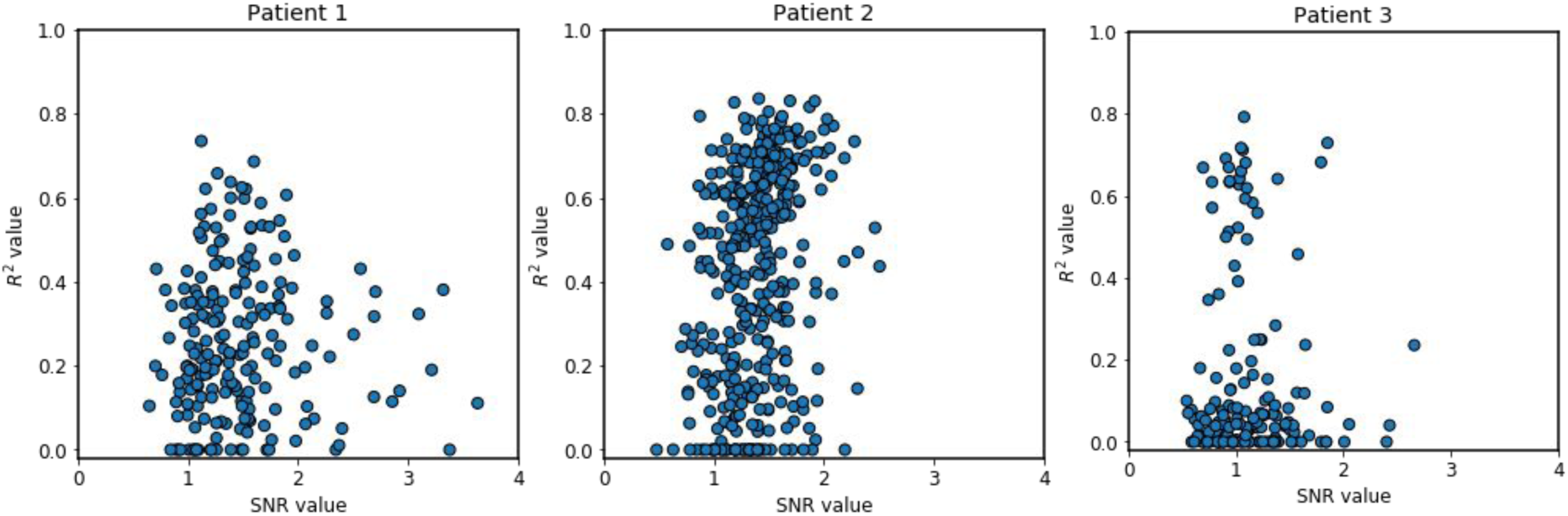
The distribution of the wave model *R*^2^ as a function of the SNR in the individual spikes for the three patients analyzed. The scatter plots do not exhibit any significant dependence of the goodness of fit metrics (*R*^2^) and the SNR of individual spikes.

To scrutinize this result further, we analyzed the distribution of *R*^2^ for each cluster separately (Figure 10). None of the clusters showed a pronounced dependence of the goodness of fit on the SNR. Interestingly, the clusters with the largest proportion of wave-like spikes (framed in red) show a separate cloud of dots with large *R*^2^ values located above the same SNR range as the dots with low *R*^2^.

**Figure 10:**
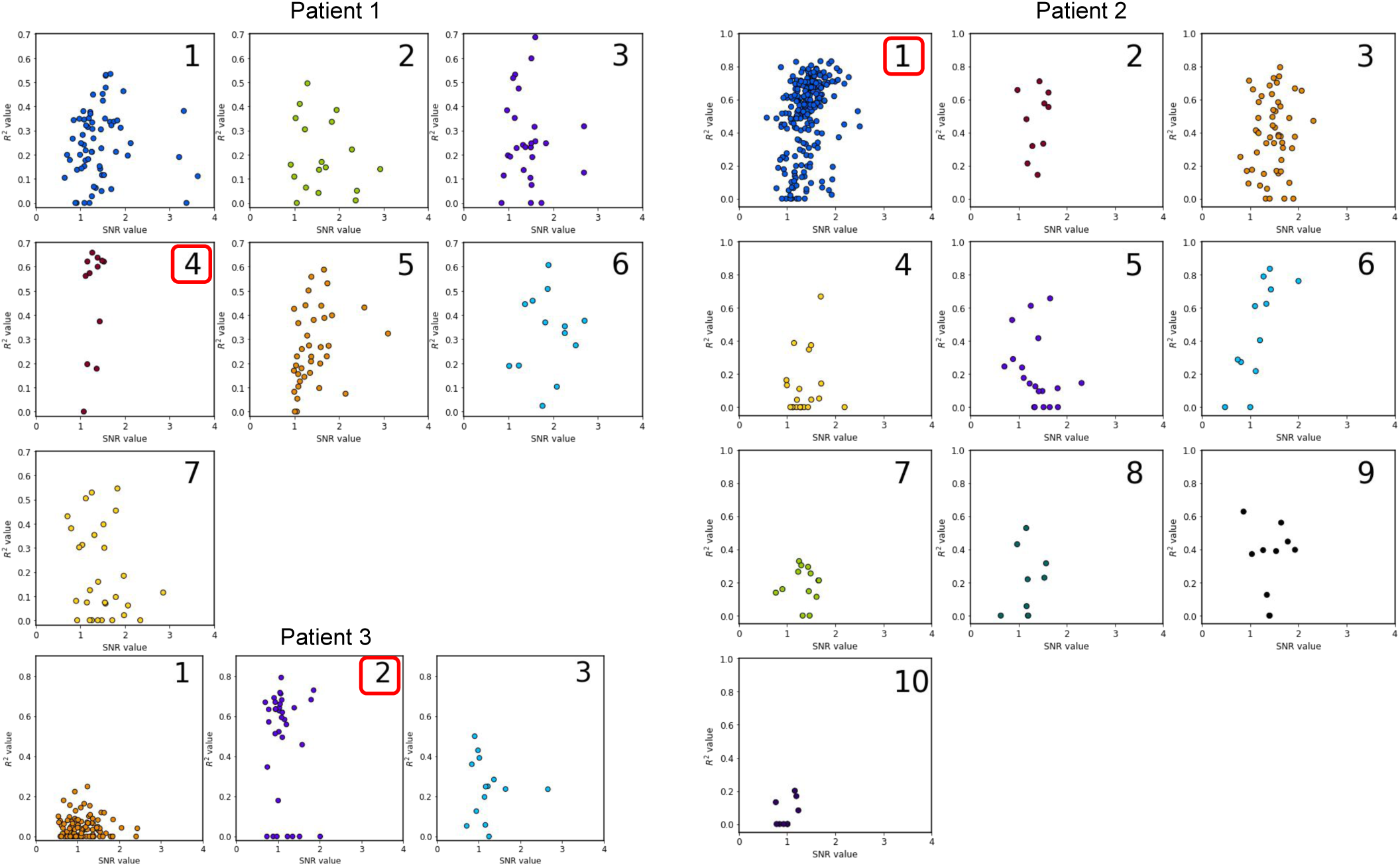
The distribution of the wave model *R*^2^ as a function of the SNR in the individual spikes for each of the clusters in the three patients. For each patient the red boxes frame the index of the cluster with the largest percentage of wave-like spikes. Non of the clusters exhibits a significant dependence of the goodness of fit metrics (*R*^2^) on the SNR of individual spikes.

In all three datasets analyzed, the clusters vary by the percentage of spikes well-explained by the traveling wave model. Intriguingly, the regions with the highest percentage of propagating spikes for Patient 1 and Patient 2 coincide with the epileptogenic focus determined independently by the neurosurgeons and confirmed by Engel I outcome as determined by the two-year follow-up observation of the patients. The information about the location of the epileptogenic region in Patient 3 is unavailable because no surgery was performed. These findings are in-line with the previously reported observation of consistent propagating patterns in the epileptogenic region [33].

## 4 Discussion

The novel regularization technique developed here solves the MEG inverse problem using traveling wave priors. As such, this algorithm is applicable to a range of neurophysiological inquiries regarding the mechanisms and functions of propagating activity in the brain. In this study, we applied our method to the analysis of interictal spikes, which we modeled as travelling waves. We used LASSO technique with the positively constrained coefficients to derive the optimal propagation velocity of the waves. This technique was tested on both simulated and real MEG data. We demonstrated that the dynamics of propagating MEG spikes could be measured at a millimeter/millisecond spatiotemporal scale. We have also observed that in all the three patients analyzed wave-like behavior was characteristic only for the spikes from a single well defined cortical region.

While the proposed method successfully identified spike travelling waves and reconstructed their anatomical paths, it was still prone to errors related to (1) the uncertainties in the estimation of the wave starting point, and (2) inaccuracies of cortical surface parametrization. The first source of errors can be ameliorated by selecting high-amplitude spikes for the analysis. The second issue can be dealt with by performing brain scans with a 7-T MRI machine that maps brain anatomy more precisely.

Despite these issues with errors, a fraction of interictal spikes was described well by the travelling-wave representation. Moreover, these spikes originated from a single cortical location and for the patients in whom SOZ data were available these clusters of spikes corresponded to the SOZ. Based on these findings that align well with invasive findings [30, 6], we suggest that the travelling wave analysis of interictal MEG data could aid the localization of SOZ. Yet, not all spikes could be explained by the wave propagation model with a small number of dominant propagation directions. These cases will need to be studied in more detail in the future.

At this point, we conclude that representation of interictal spikes as travelling waves describes well at least part of the data from epileptic patients, which indicates that epileptic brain activity is a wave phenomenon. The practical implication of this finding is that in the future the analysis of local propagation patterns of interictal activity may become an essential part of pre-surgery diagnostics and contribute to a differentiation of epilepsy types, EZ localization and planning of a maximally sparing surgical strategy to resect the epileptogenic tissue. The algorithm used to track the local spatial-temporal dynamics of interictal spikes can be directly extended to analyze seizure onset segments which in the nearest future may become less problematic with wearable sensor arrays such as those based on optically pumped magnetometers.

We foresee that the proposed approach for the analysis of epileptic activity and solving the MEG inverse problem will become more accurate and beneficial for the patients in the future, when advanced MEG instrumentation will emerge. As such, our computational method could become a valuable tool for quantitatively advanced, noninvasive diagnostics of patients suffering from epilepsy and for precise pre-surgical mapping of epileptogenic regions in patients with drug-resistant epilepsy.

